# Mapping Protein Occupancy on DNA with an Unnatural Cytosine Modification in Bio-orthogonal Contexts

**DOI:** 10.1101/2025.11.24.690163

**Authors:** Ruiyao Zhu, Christian E. Loo, Christina M. Hurley, Jared B. Parker, Rahul M. Kohli

## Abstract

The epigenome provides a dynamic layer of gene regulatory control above the static genetic sequence. DNA base modifications are key epigenetic regulators, predominantly found within CpG contexts in mammalian genomes. Working in tandem with these DNA modifications, chromatin-associated proteins and transcription factors further control gene expression. Given the interplay of these factors, concurrent mapping of DNA base modifications with protein-DNA occupancy can greatly aid in interpreting the epigenome. Existing multi-modal mapping methods include the use of DNA methyltransferases to mark accessible, protein-unbound DNA in non-CpG contexts. However, such approaches can either confound readouts with native DNA modifications or constrain users to third-generation sequencing approaches. To circumvent these limitations, we explored the possibility of introducing an unnatural DNA base modification, 5-carboxymethylcytosine, as an alternative label for protein occupancy. Here, we report our efforts to rationally engineer non-CpG-specific DNA methyltransferases to take on neomorphic DNA carboxymethyltransferase (CxMTase) activities. We find that DNA carboxymethylation of cytosines in GpC contexts shows broad compatibility with the most widely used epigenetic detection methods and can be used to reliably report on protein occupancy states. Using this approach, we reveal the single-molecule binding patterns of LexA, a master repressor in the bacterial DNA damage (SOS) response, at its self-regulated and endogenously methylated promoter. We thus show that unnatural DNA modifications in bio-orthogonal contexts can uncover biological insights and potentiate new approaches to multimodal epigenetic profiling.

**TOC GRAPHIC:** 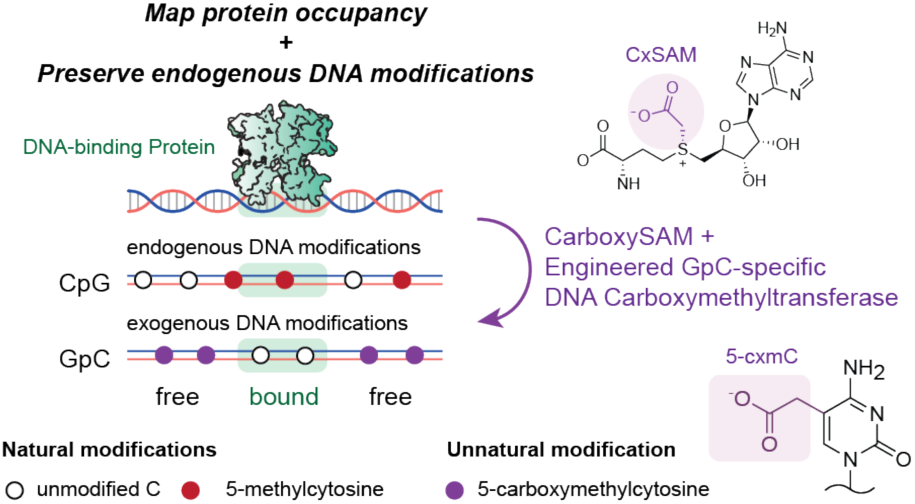

## INTRODUCTION

Chemical modifications of DNA bases enable dynamic control over gene regulation and cell fate.^1^ In mammalian genomes, DNA base modifications primarily occur at cytosines within cytosine–guanine (CpG) dinucleotide contexts, with 5-methylcytosine (5mC) being the most common.^2^ 5mC is generated by DNA methyltransferases (MTases) using *S*-adenosyl-L-methionine (SAM) as a methyl donor.^3^ 5mC is typically regarded as a repressive epigenetic mark with diverse roles including transcriptional silencing, imprinting, and X-chromosome inactivation.^2^ 5mC dynamics are regulated in part by ten–eleven translocation (TET) dioxygenases, which can oxidize 5mC to 5-hydroxymethylcytosine (5hmC) as a step towards relieving 5mC-associated gene repression.^4^ Importantly, the functional consequences of these DNA base modifications depend critically on their context, whether buried within nucleosome-wrapped DNA or exposed and available to interact with transcription factors (TFs), including repressors and activators.^5,6^ In distinction to mammalian genomes, methylation plays well established roles in discrimination of self versus non-self in bacteria, with relatively more underexplored effects on gene regulation and TF binding.^7^

Given the interconnected nature of the regulatory landscape in both eukaryotes and prokaryotes, jointly profiling DNA modification and protein occupancy is essential. Numerous methods have been established to profile individual layers of regulatory networks, localizing either DNA modifications or protein occupancy.^8,9^ Conventional chemical approaches can be used to map these layers, including methods such as bisulfite sequencing (BS-Seq) to discriminate cytosine modification states or hydroxyl radical footprinting to profile protein-unbound regions.^10,11^ However, chemical approaches can be destructive, requiring large DNA inputs and causing biased DNA fragmentation.^12,13^ To overcome these limitations, alternative approaches have recently harnessed DNA modifying enzymes. For example, APOBEC3A (A3A), a DNA cytosine deaminase involved in viral defense, has been repurposed to profile DNA cytosine modifications in methods such as APOBEC-coupled epigenetic sequencing (ACE-Seq).^14,15^ Other enzymatic approaches, such as DNase-Seq and ATAC-Seq, have been used to fragment and thereby map open chromatin.^16,17^ In contrast to chemical approaches, these enzymatic approaches are non-destructive, enabling the analysis of rare cell populations or even single cells.

For protein-occupancy mapping, DNA fragmentation can limit resolution across long reads and be associated with biases.^18,19^ Consequently, enzymatic approaches that selectively label protein-unbound DNA with DNA modifications have also been developed. Methods like SMAC-Seq, SAMOSA, and Fiber-Seq use adenine DNA MTases to mark accessible regions with 6-methyladenine (m6A), a modification absent in mammalian genomes.^20–22^ Other approaches including NOMe-Seq and its variants instead leverage a GpC-specific cytosine MTase M.CviPI to selectively install 5mC in DNA regions unbound by proteins. The resulting exogenous 5mC can be partially distinguished based on sequence context, allowing for simultaneous profiling of DNA methylation and accessibility.^23–25^ Despite these advances, the use of natural modifications as protein occupancy labels presents inherent limitations. In m6A-based methods, the requirement for third-generation sequencing to read m6A significantly increases sample input requirements and limits the use of established methods, such as BS-Seq, for profiling cytosine methylation.^26^ With 5mC-based methods, resolution of 5mC from 5hmC and disambiguation of 5mC in overlapping CpG and GpC contexts are key limitations.

Here, we considered the possibility that the enzymatic introduction of *unnatural* base modifications could complement current multimodal mapping methods. In support of this concept, we recently showed that a CpG-specific DNA MTase (M.MpeI) can be engineered to gain a neomorphic ability to use carboxy-*S*-adenosyl-L-methionine (CxSAM) as a substrate.^27^ The engineered M.MpeI N374K uses CxSAM to convert unmodified CpGs into ones with 5-carboxymethylcytosine (5cxmC), an unnatural base modification that resists both bisulfite and enzymatic deamination. This CpG-specific DNA carboxymethyltransferase (CxMTase) enabled the development of Direct Methylation sequencing (DM-Seq), a fully enzymatic method for directly profiling 5mC alone at single-base resolution.^28^

The precedent established by the CpG-specific DNA CxMTase suggests the possibility that unnatural DNA modifications could also be used as orthogonal labels in non-CpG contexts to map DNA-protein interactions (**Figure 1**). Here, we engineer and characterize novel CxMTases with non-CpG specificities. We further demonstrate their ability to act as sensors of protein occupancy, allowing us to newly map the binding pattern of LexA, the key TF controlling the bacterial DNA damage response, on differentially methylated target sequences. We thus show that engineered CxMTases can support a fully enzymatic pipeline for mapping of DNA-protein interactions, extending the utility of DNA modifying enzymes in resolving heterogeneous epigenetic states.

**Figure 1.**
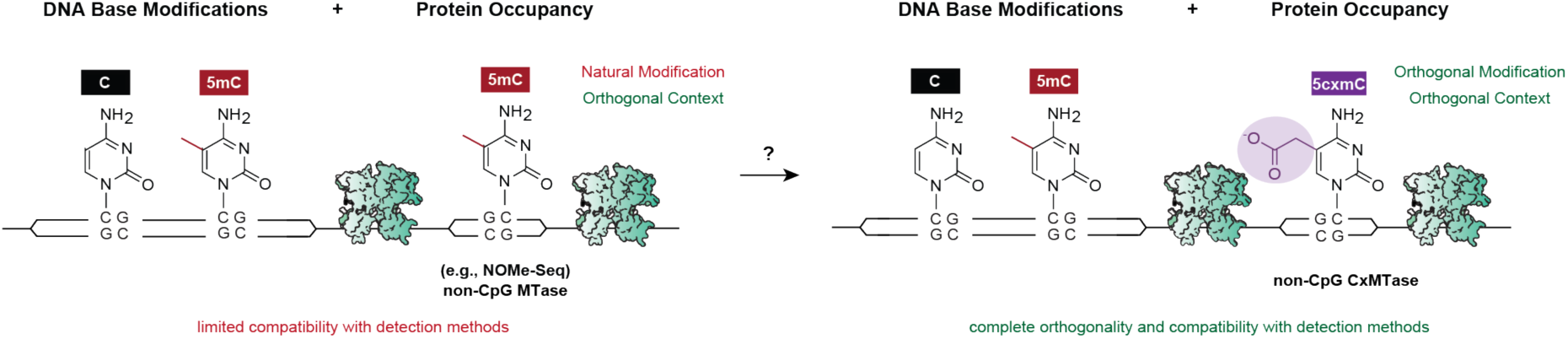
Unnatural modifications in non-native contexts for joint profiling of epigenetic bases and protein occupancy. Natural DNA base modifications in mammalian genomes include 5mC (red) in CpGs. 5mC in non-CpG contexts has been used to label protein-unbound DNA in methods like NOMe-Seq (left). The chemical identity overlap limits the compatibility with different detection methods. 5cxmC (purple) is an unnatural DNA base modification resistant to chemical and enzymatic conversion. Explored here is the concept that 5cxmC installed in non-CpG contexts (right) could be used for protein occupancy mapping to enable wider compatibility.

## RESULTS AND DISCUSSION

### Construction and Screening of Non-CpG-specific MTase Candidates

In exploring candidates to introduce unnatural labels onto DNA, we considered that avoiding overlap with endogenous cytosine modifications in CpGs could offer long-term opportunities to study mammalian DNA modifications. Additionally, as the frequency of the MTase recognition sequence determines the resolution of protein occupancy footprinting of DNA, shorter recognition sequences could offer higher resolution. With both of these considerations in mind, we focused on two previously identified cytosine DNA MTases that each have a specific dinucleotide recognition sequence: M.CviPI (Gp<u>C</u>) and M.CviQIX (<u>C</u>pC) (target cytosine underlined).^29^

Given the conserved nature of MTases, we hypothesized that mutations at residues homologous to M.MpeI N374 could potentially confer carboxymethylation activity in other DNA cytosine MTases.^30^ As homologous residues were not immediately obvious from a primary sequence alignment, we generated AlphaFold3-predicted structures of M.CviPI and M.CviQIX bound to their respective DNA substrates,^31,32^ and aligned to the DNA-bound structure of M.MpeI (PDB 4DKJ).^33^ The aligned structures showed high similarity around the active site, exhibiting features of a Rossmann fold and the base-flipping mechanism common to DNA MTases (**Figure S1**). Using these models, we identified M.CviPI Y205 and M.CviQIX N218 as the residues closest to the Cα of the N374 site in M.MpeI and therefore targets for potential mutagenesis (**Figure 2**). In exploring N374 of M.MpeI with saturation mutagenesis, we had found that the introduction of a cationic side chain best supported CxSAM activity, which could be rationalized in molecular modeling by the establishment of a new salt-bridge interaction.^27,34^ Building on this precedent, we therefore chose to focus our efforts on K and R mutants targeting Y205 in M.CviPI and N218 in M.CviQIX.

**Figure 2.**
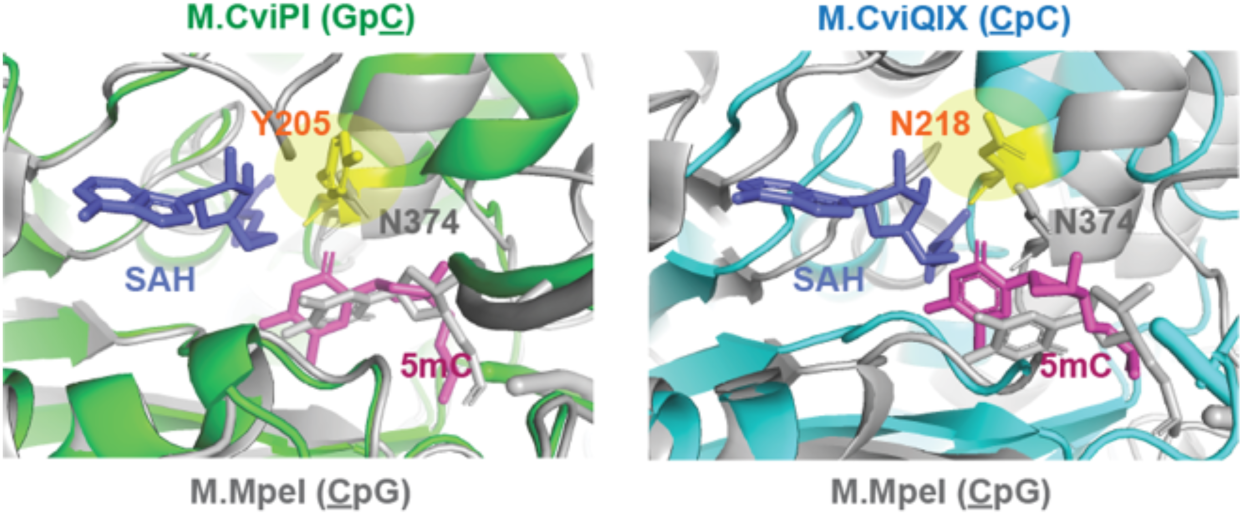
Homologous residue targets for mutagenesis to confer carboxymethylation. AlphaFold3-predicted structures of M.CviPI (left, green) and M.CviQIX (right, cyan) aligned with the crystal structure of M.MpeI (grey, PDB 4DKJ) bound with DNA with a 5mC (pink) and the side product SAH (dark blue). Target residues, Y205 in M.CviPI and N218 in M.CviQIX, are highlighted in yellow.

Given that high enzyme activity is essential for method development, we first evaluated the impact of potential mutations on canonical MTase activity, recognizing that prior work on M.MpeI suggested that gain in CxMTase activity was accompanied by maintained MTase activity.^27^ To this end, we individually expressed the wild-type (WT) and variant enzymes in a methylation-tolerant strain of *E. coli*, using WT M.MpeI as a comparison. We then assessed MTase activity on host DNA *in vivo* by extracting genomic DNA (gDNA) from the *E. coli* and digesting it with HaeIII, a GG<u>C</u>C-recognizing restriction enzyme blocked by modification at the internal cytosine (**Figure 3A**). As expected, WT M.MpeI did not protect against HaeIII digestion, consistent with CpG-specific methylation. In contrast, both WT M.CviPI and M.CviQIX conferred significant protection, indicating that most GG<u>C</u>C sites in the gDNA were modified. With both K and R mutants for each enzyme, we observed a shearing pattern consistent with partial protection, suggesting reduced but detectable activity. Notably, the M.CviPI mutants exhibited higher levels of protection relative to M.CviQIX (**Figure 3B**), suggesting higher MTase activity in this setting. We therefore proceeded with the GpC-specific M.CviPI for more detailed biochemical investigation.

**Figure 3.**
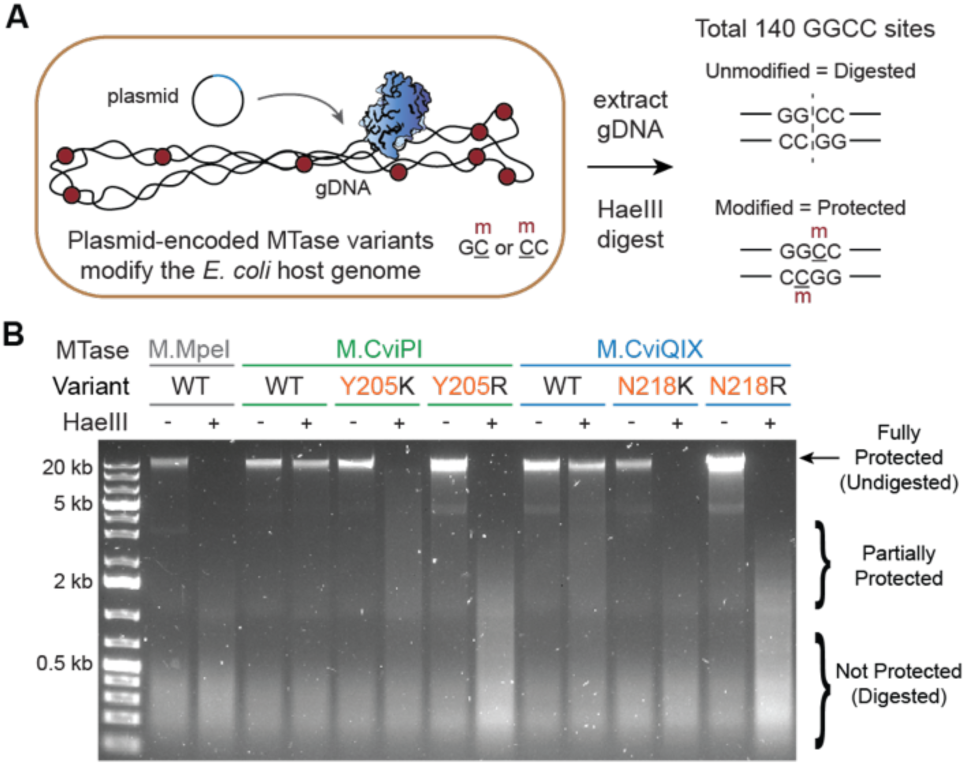
*In vivo* activity screen of MTase candidates. (A) Scheme of activity screen in *E. coli*. Genomic DNA (gDNA) is extracted from *E. coli* expressing the plasmid-encoded MTase variants. GG<u>C</u>C sites in the genome can be modified by active M.CviPI (Gp<u>C</u>) or M.CviQIX (<u>C</u>pC) using intracellular SAM as a co-factor, while M.MpeI (<u>C</u>pG) is not expected to modify this site. Modified DNA is protected from restriction digest by HaeIII. (B) Agarose gel showing undigested or HaeIII digested gDNA recovered from cells expressing different MTase variants.

### Engineered MTases Gain Neomorphic Carboxymethylation Activity

Given that SAM is 400-fold more abundant than CxSAM in *E. coli*, we attributed the activity detected upon enzyme overexpression predominantly to SAM-mediated generation of 5mC.^35^ To examine enzymatic activity with SAM and CxSAM *in vitro*, we next expressed and purified M.CviPI WT, Y205K, and Y205R (**Figure S2**). We then incubated an unmodified pUC19 plasmid DNA substrate with increasing concentrations of each enzyme in the presence of either SAM or CxSAM. Modification levels were again assessed by HaeIII digestion (**Figure 4A**). As anticipated, pUC19 DNA incubated with WT M.CviPI was efficiently digested by HaeIII in the absence of SAM, but protected in its presence, consistent with efficient GpC methylation. In line with the *in vivo* screen, both Y205K and Y205R mutants also protected pUC19 from digestion in the presence of SAM. However, unlike the *in vivo* screen examining gDNA protection, the methylation efficiency appeared similar between WT and mutants *in vitro* (**Figure 4B**). This discrepancy likely reflects limiting SAM concentrations *in vivo*, which disproportionately affected mutants. When incubated with CxSAM, WT M.CviPI failed to protect pUC19 from HaeIII digestion, consistent with its constrained active site and inability to utilize CxSAM. By contrast, both Y205R and Y205K incubated with CxSAM protected pUC19 from HaeIII digestion, supporting their efficient use of CxSAM as a cofactor and our hypothesis.

**Figure 4.**
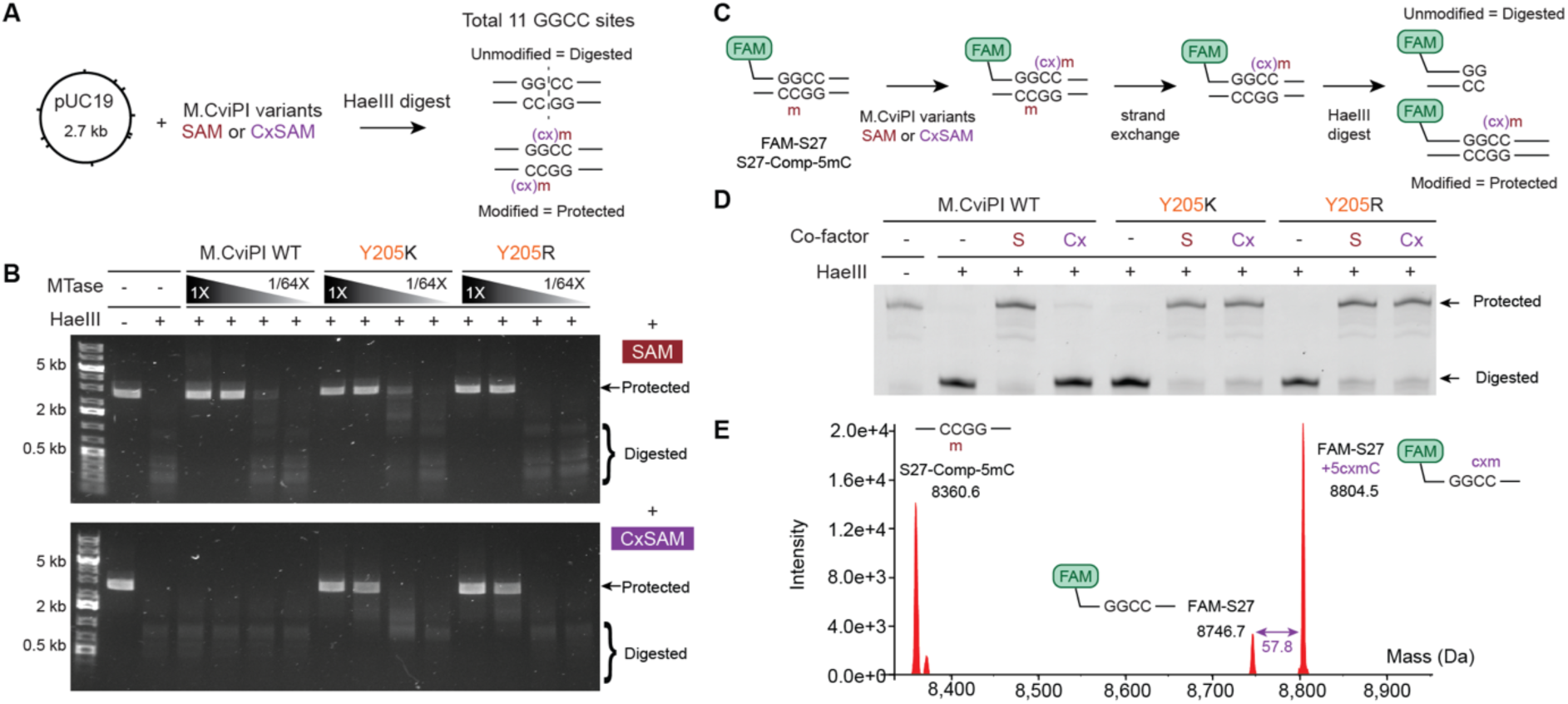
*In vitro* activity assays of MTase candidates. (A) Scheme of pUC19-based activity assay. The unmodified pUC19 substrate is incubated with M.CviPI variants and either SAM or CxSAM. The substrate is then purified and digested by HaeIII to report on modification status. (B) Agarose gel of restriction digest results. Modification is detected as protection from HaeIII digestion. Enzyme variant concentrations were normalized and titrated with 4-fold serial dilutions from 1X to 1/64X while SAM/CxSAM concentration is fixed at 300 µM. (C) Scheme of oligonucleotide-based activity assay. The FAM labeled top strand (FAM-S27) containing a single GG<u>C</u>C site is annealed with a methylated complement. The duplex was incubated with M.CviPI variants and SAM or CxSAM. The methylated bottom strand is exchanged for an unmodified strand to permit evaluation of protection with HaeIII digestion. (D) DNA Denaturing PAGE of duplexes analyzed by the activity assay where each M.CviPI variant was incubated with no (-), 300 µM SAM (S), or 300 µM CxSAM (Cx). (E) Intact LC/MS trace of oligonucleotide duplex treated with M.CviPI Y205K and CxSAM that displays the expected mass shift for the carboxymethylated oligonucleotide.

To further characterize the potential CxMTase activity, we next performed an orthogonal assay using a duplexed oligonucleotide, rather than plasmid DNA, as the substrate. In this assay, we used a fluorescently labeled 27-bp oligonucleotide (FAM-S27, **Table S1**) with a single GpC site within an HaeIII recognition sequence. In the unlabeled complement strand, we included 5mC at the GpC site (S27-Comp-5mC), allowing us to focus on activity at the single unmodified GpC in the labeled strand without confounding by the opposite strand. Following incubation with the M.CviPI variants and either SAM or CxSAM, the complementary strand was exchanged with one lacking 5mC (S27-Comp-C) and products interrogated with HaeIII to determine whether the GpC in FAM-S27 is unmodified (digested) or modified (protected from digestion) (**Figure 4C**). In this assay, we observed that both WT and mutant enzymes incubated with SAM protected FAM-S27 from HaeIII digestion, consistent with retention of MTase activity. By contrast, with CxSAM, only the Y205K and Y205R mutants showed significant protection, consistent with a neomorphic gain of CxMTase activity (**Figure 4D**). To definitively establish the modification identity, we performed intact oligonucleotide LC/MS on the initial duplex product. In addition to the two expected peaks corresponding to the substrate, we observed a unique peak with a mass shift of 57.8 Da relative to FAM-S27, consistent with a single 5cxmC introduced in the product oligonucleotide (**Figure 4E**). Thus, the rationally engineered M.CviPI variants show definitive CxMTase activity, positioning them for further applications.

### Bio-orthogonal Carboxymethylation Offers Compatibility with Common Deamination Methods

Having validated successful GpC-specific CxMTase engineering, we next aimed to quantitatively characterize activities on a more complex DNA substrate and explore compatibility with different epigenetic sequencing methods. To that end, we performed transfer reactions on an unmodified λ phage gDNA (48.5 kb) with either SAM or CxSAM, and then resolved the resulting DNA modifications at base resolution (**Figure 5A**). The modified DNA was sheared and sequenced to >100-fold depth using both BS-Seq and ACE-Seq. The use of both chemical and enzymatic sequencing approaches allows for resolution of C, 5mC, and 5cxmC. Based on established precedent, both 5mC and 5cxmC are protected from C-to-T conversion in BS-Seq, whereas only 5cxmC is protected from C-to-T conversion in the ACE-Seq pipeline.^28^ As a positive control, we also included the engineered M.MpeI N374K CpG-specific CxMTase to allow for comparison of CpG- and GpC-specific MTase activities.

**Figure 5.**
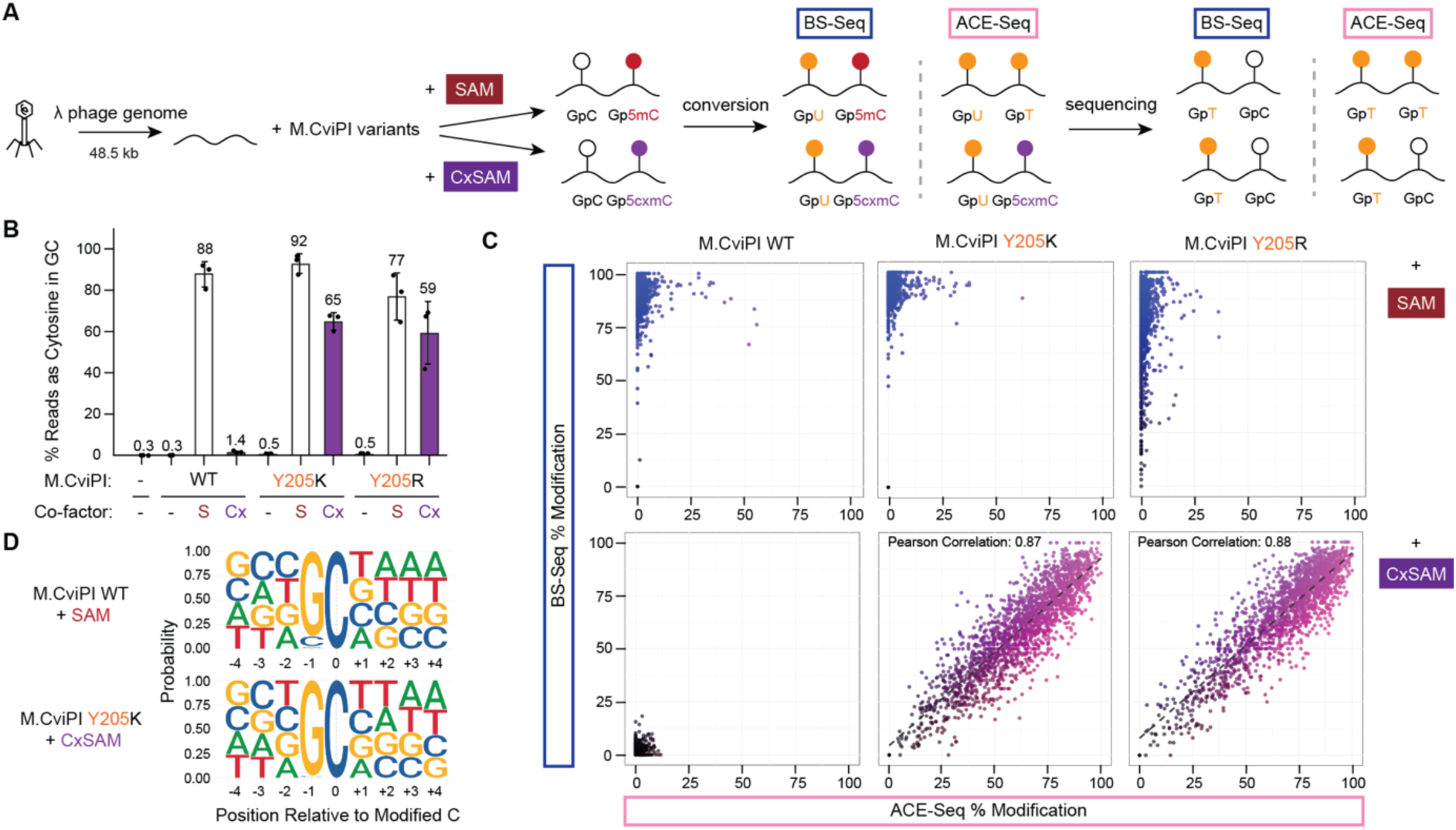
Base-resolution characterization of modification patterns on λ phage genomic DNA. (A) Scheme of sequencing-based characterization. (B) Cytosine modification is measured by protection of GpC sites from C-to-T conversion (% reads as C) in GpC contexts. Shown are individual data points from independent replicates (n = 3), with mean values in bars and standard deviation noted. (C) Correlation plots between BS-Seq and ACE-Seq after reaction of λ phage gDNA with different M.CviPI and co-factor pairs. Each dot represents a single GpC site in the gDNA. Data is filtered for GpC sites with >15 total reads and randomly downsampled to 2000 sites to avoid overplotting. A strong Pearson correlation is indicated by a dashed line in the last two subplots. (D) Modification logos of different enzyme and co-factor pairs. The numbering is relative to the target cytosine base, which is the 0 position.

First, focusing on BS-Seq, we observed the efficient use of SAM across enzyme variants as measured by the average level of protection of GpC sites from C-to-T conversion (**Figure 5B**). This result suggests efficient MTase activity across various GpC contexts, consistent with our previous *in vitro* assays that only interrogated GGCC sites using HaeIII cleavage. At the SAM concentration examined, there was no significant difference in global modification levels between M.CviPI WT and mutants. Using CxSAM with the WT enzyme, we observed no significant protection of GpC sites from C-to-T conversion relative to background. By contrast, when the Y205K and Y205R mutants were used with CxSAM, average protection levels of 64% ± 4% and 59 ± 15% were observed. Notably, all M.CviPI variants showed no significant modification in WCG contexts as expected (**Figure S3A**).

Unlike 5mC generated from SAM, 5cxmC generated from CxSAM is known to resist enzymatic deamination in the ACE-Seq pipeline, opening up the possibility that 5cxmC mapping could be done with all-enzymatic approaches.^28^ Indeed, when we focused our analysis on individual GpC sites in the genome when SAM was used as a substrate, the sites unconverted in BS-Seq now show significant C-to-T conversion in ACE-Seq, leaving no readily detectable signal. By contrast, samples treated with the M.CviPI mutants and CxSAM yielded unconverted GpC sites in both BS-Seq and ACE-Seq. Moreover, these signals from ACE-Seq correlate with those from BS-Seq (**Figure 5C**), confirming that 5cxmC protects the GpC sites from both chemical and enzymatic conversion, enabling compatibility with multiple detection methods.

To further characterize context specificity, we generated a position weight matrix of the modified sites under various conditions (**Figure 5D**; **Figure S3B**; **Figure S3C**). As expected, all variants show a GpC context preference. WT M.CviPI is known to have some off-target activity at CpC sites, which can confound footprinting in NOMe-Seq.^23^ Interestingly, while our BS-Seq revealed this expected off-target CpC activity with WT M.CviPI and SAM, context specificity improved for both mutants without compromising overall activity. These observations suggest that M.CviPI Y205K and Y205R, even when used in conjunction with SAM, may offer useful advantages. With the mutant enzymes, the selectivity pattern is yet further enhanced with CxSAM, with minimal activity detectable outside of a GpC context. We also examined the sequence context flanking target sites (**Figure S4**) to look for bias in GpC modification. We observed that the CpG-specific CxMTase shows higher bias, with discrimination against −1 A or +2 R (A or G) relative to the CpG site, while the GpC-specific CxMTase shows lower overall bias, with only minor discrimination against +1 A relative to the GpC site.

In prior work, we found that the CpG-specific CxMTase M.MpeI N374K favors the generation of hemi-carboxymethylated CpG sites, likely due to the steric bulk of the first carboxymethyl group.^28^ This asymmetric activity posed a challenge in the development of DM-Seq. To similarly examine GpC-specific CxMTases, we correlated the modification on the Watson strand with those in the Crick strand for GpC dyads across the λ phage genome (**Figure S3D**; **Figure S3E**). Compared to the CpG CxMTase, the GpC CxMTases show significantly less asymmetry, as evidenced by the limited number of sites where modification on one GpC is correlated with lower levels on the opposite GpC. Additionally, while CpG CxMTase asymmetry limits global modification to approximately 50%, GpC CxMTases do not exhibit the same constraint. Indeed, by increasing the CxSAM concentration from 300 μM to 800 μM carboxymethylation can be enhanced to 82.7% of sites without notable asymmetry or off-target activity (**Figure S3C**). Thus, GpC CxMTases could offer some distinctive advantages over CpG CxMTase by supporting the potential symmetric introduction of 5cxmC on both DNA strands.

### GpC Carboxymethylation Maps TF Binding and DNA Modifications on Single Molecule

Having engineered GpC CxMTases, we last aimed to apply the neomorphic enzyme as a potential reporter of protein-bound states to offer new biological insights. We chose to focus on regulatory control of the bacterial DNA damage response, also known as the SOS response.^36^ LexA is the master SOS transcriptional regulator that, in the absence of DNA damage, binds to and represses a large collection of DNA damage response genes through specific interactions with DNA binding motifs in the gene promoters. These SOS boxes are characterized by two conserved 5′-CTG(A)-3′ motifs separate by 10 bps with variable intervening sequence that modulates LexA affinity.^37^ In the presence of DNA damage, LexA is triggered to undergo self-proteolysis, resulting in relief of gene repression and induction of the SOS response. Notably, LexA expression is self-regulated with a distinctive *lexA* promoter structure including two juxtaposed SOS boxes, one of which contains an embedded site for endogenous Dcm methylation (C<u>C</u>WGG).^38^ While previous studies have examined each of the SOS boxes separately in synthetic oligos, it remains unknown if both SOS boxes can be occupied simultaneously and whether occupancy on this native dual SOS box promoter could be altered by methylation (**Figure 6A**).^39^

**Figure 6.**
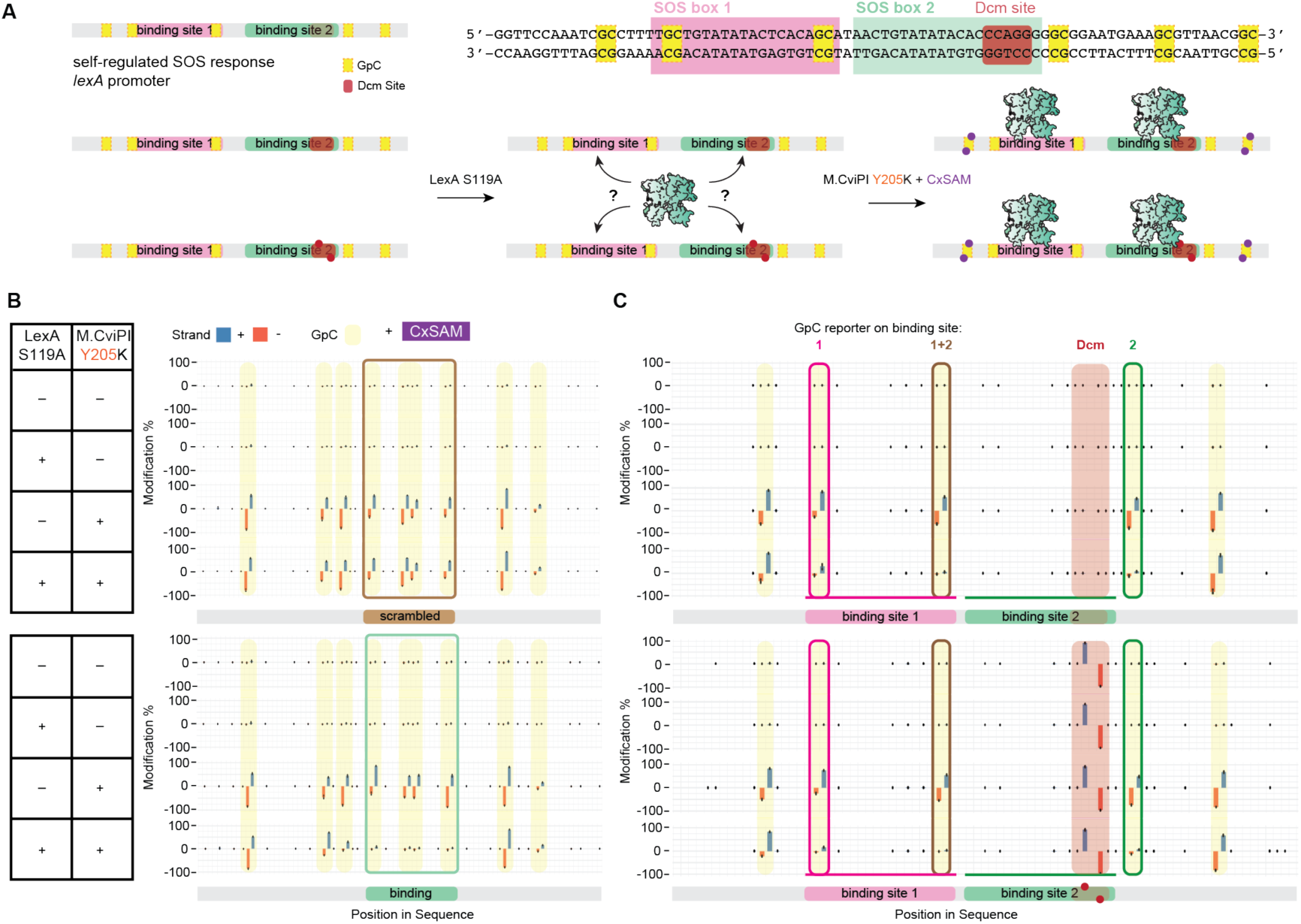
CxMTase footprinting assay for LexA-operator binding. (A) At top, schematic of the *lexA* promoter, with two SOS binding boxes (pink/green) and an endogenous methylation site (red). At bottom, schematic of the LexA-operator footprinting assay on unmethylated or methylated DNA (red circles), denoting the possible 5cxmC additions (purple circles). (B) Footprinting validation for LexA binding at substrates containing either a single scrambled or unscrambled SOS box. Modification level from 0 to 100% are noted as assessed by BS-Seq, with + strand (blue, upward) and – strand (orange, downward). Mean values are shown (n = 3) as bars with standard deviation. The scrambled 20-bp LexA binding site is indicated by the brown colored box. The 20-bp LexA binding site is indicated by the green colored box. (C) Footprinting assay at LexA self-regulated operator. Specific GpCs are noted as potential reporters on binding at Site 1, Site 2, or both Site 1+2.

Before attempting to dissect the mechanism of *lexA* self-regulation, we first assessed whether the GpC-specific CxMTase can report protein-bound states. In established NOMe-Seq approaches, WT M.CviPI selectively introduces 5mC modifications into DNA unencumbered by DNA-binding proteins that block MTase activity. To model whether the bio-orthogonal 5cxmC mark could similarly be used, we devised an *in vitro* assay to assess functionality. In our assay design, we employed a LexA S119A mutant capable of binding to DNA but unable to self-proteolyze. We designed an oligonucleotide containing an embedded central SOS box, which included GpC dinucleotides in a HhaI cleavage site (GCGC), along with a matched control where the SOS box motif was scrambled (**Table S1; Figure S5A**). Using duplexes amplified with a 5′-fluorescent label, we validated that LexA S119A binds to the SOS-box-containing duplex, but not the scrambled duplex (**Figure S5B**).

Using these two duplexes, we next permutated two added factors: 1) incubation in the presence or absence of LexA S119A and 2) subsequent incubation with either the SAM/WT M.CviPI or the CxSAM/M.CviPI Y205K substrate/enzyme pair. As an initial assessment, the resulting duplexes were purified and subjected to methylation-sensitive HhaI digestion to interrogate the modification state of the target GCGC site within the SOS box (**Figure S5C**). As anticipated, in the absence of M.CviPI variants, both duplexes can be efficiently digested. In the absence of LexA, both enzyme/substrate pairs result in duplex protection, consistent with efficient modification. In the presence of LexA S119A, however, while the scrambled duplex remains protected from HhaI cleavage, the SOS box containing duplex is no longer efficiently protected, suggesting that the LexA-bound DNA is not accessible to the enzyme (**Figure S5D**).

To more rigorously examine LexA footprinting, we next used BS-Seq to resolve LexA positioning (**Figure 6B**; **Figure S6**). With the scrambled duplex, we observed that both enzyme/substrate pairs result in modifications across all GpC sites, independent of the presence or absence of LexA S119A. In contrast, with the SOS box containing duplex, we observed a significantly reduced GpC modification in the presence of LexA S119A. Importantly, this reduction was localized to the GpCs at or within +/-5 bps of the LexA binding sites, while GpCs outside of this detection window maintained similar modification patterns. Upon closer examination, we also observe CpC modification with the SAM/M.CviPI WT pair but not with the CxSAM/M.CviPI Y205K pair, consistent with our earlier evidence that the WT MTase is less selective than the engineered CxMTase. Taken together, these results indicate that the GpC-specific CxMTase can be blocked by the presence of protein bound to DNA, while reliably reporting on neighboring free DNA.

Having validated the GpC CxMTase as a reliable reporter, we turned our attention to the mechanism of *lexA* promoter self-regulation. To capture this region, we utilized a duplex containing the native 81-bp region encompassing the *E. coli lexA* promoter, and probed LexA occupancy with GpC modification using the CxSAM/M.CviPI Y205K pair (**Figure 6C**). Using this approach, we found that the presence of LexA S119A results in significantly reduced GpC modification at or within +/− 5 bps of both SOS boxes, with no significant difference between the two sites. Given the single-molecule nature of such footprinting experiments, this finding newly indicates that LexA can concurrently bind to both SOS boxes in the *lexA* promoter. We next repeated the experiment using DNA fully methylated at the embedded Dcm site.^39^ Using this substrate, we again found evidence for complete occupancy of both LexA binding sites. Additionally, consistent with a study that looked at the second SOS box in isolation,^39^ we provide single-molecule level evidence that Dcm methylation does not alter *lexA* promoter regulation.

## CONCLUSION

DNA modifying enzymes, traditionally viewed as ‘writers’, are increasingly being harnessed as ‘readers’ of the epigenetic state of DNA. Beyond resolving DNA modification states, these tools have also more recently been employed to discriminate between protein-bound vs. protein-free DNA regions, revealing insights into the multi-modal epigenetic code. While such approaches have typically relied upon the use of natural DNA MTases, we sought to develop a bio-orthogonal label to report on protein occupancy, enabling broad compatibility with common epigenetic sequencing methods and more versatile readouts.

Here, we employed structure-guided mutagenesis to MTases that act in non-CpG contexts to yield new CxMTases. We showed that the GpC-specific CxMTase can efficiently introduce 5cxmC, a mark compatible with both chemical (BS-Seq) and enzymatic (ACE-Seq) approaches for mapping DNA modifications. Further, we determined that 5cxmC can be used to mark non-protein-bound DNA. This work builds on our prior success engineering the CpG-specific CxMTase M.MpeI N374K, showcasing that similar mutations at structurally homologous residues within M.CviPI can also result in CxMTase activity. Our findings support a generalizable strategy for converting MTases to CxMTases and suggest that this residue, originally thought to play a key role in MTase activity, is permissive to variation, offering a means to expand cofactor selectivity.^40,41^ Complementing prior studies that have focused on ‘space opening’ mutations that permit use of bulky SAM analogs, we show that introducing cationic residues can enhance catalysis with CxSAM. These results demonstrate the potential of introducing new molecular features, such as charge-to-charge interactions, in targeted partnerships between engineered MTases and SAM analogs.^42,43^

While generating the neomorphic CxMTase highlights some shared plasticity in MTases, the GpC-specific CxMTase variants we characterized have some distinctive attributes that make them particularly useful tools. For one, in contrast to our prior CpG CxMTase, we found that the GpC CxMTase activity was not affected by carboxymethylation on the opposite strand. Further study could elucidate the mechanistic determinants of the symmetric activity of CxMTases, which help overcome limitations with sequencing methods such as DM-Seq. Second, while the WT M.CviPI is known to have some non-GpC off-target activity, the M.CviPI Y205K showed higher fidelity for acting on GpC sites, with either SAM or CxSAM as substrates. Thus, the engineered MTases may also offer distinct advantages in NOMe-Seq and other pipelines involving the use of M.CviPI WT. ^44–47^

Most notably, the GpC-specific CxMTases can generate unnatural modifications to serve as reporters of the protein-DNA occupancy by selectively labeling unbound DNA. Specifically, with the *lexA* promoter harboring two LexA binding sites and an embedded site for endogenous methylation, footprinting offers the single molecule resolution that allows us to conclude that both sites can be occupied concurrently, independent of DNA methylation. Our findings are intriguing in that LexA binding is known to bend the DNA helix, suggesting that these biophysical features do not disrupt LexA binding at a neighboring site.^48^ Our observations suggest that self-regulation of LexA likely reflect stringent controls related to the simultaneous binding at tandem SOS boxes, but that the presence of an endogenous methylation site may only be incidental. While further work is needed to benchmark the performance of CxMTase in more complex workflows with mammalian genomic DNA samples, our work presents proof of concept for the use of unnatural DNA base modifications as a novel, non-destructive approach to footprint protein occupancy without fragmentation or interference with native DNA base modifications.

Using unnatural DNA modifications as protein occupancy markers brings unique advantages over previous approaches that rely on natural modifications. For one, 5cmxC resists both chemical and enzymatic deamination and can therefore overcome the inability of exogenous methylation labels to be retained in next generation sequencing-based workflows. For example, as both 5hmC and 5cxmC are resistant to enzymatic deamination with A3A, the GpC-specific CxMTase should be useful for multi-modal profiling of 5hmC together with open chromatin, analysis that is not readily feasible with current approaches. Moreover, the GpC-specific CxMTase opens new possibilities for long-read sequencing platforms. We posit that 5cxmC, given its distinct chemical character, will likely generate a unique signal from natural base modifications, potentiating a versatile solution for jointly mapping DNA modification and protein occupancy at the single-molecule level. It is worth noting that the line between unnatural and natural modifications remains blurry. While 5cxmC remains an unnatural DNA modification in the context of gene regulation for higher-order organisms, the modification has been recently discovered in phage genomes, suggesting that nature itself could be an inspiration for future unusual DNA modifications with utility in new sequencing technologies.^49^ More generally, our work highlights how the use of non-canonical DNA modifications can facilitate a more complete understanding of the functional interactions between the chemical marks on DNA bases and protein regulators in shaping gene expression.

## Notes

The University of Pennsylvania has patents pending for CxMTase enzymes with R.M.K as an inventor. C.E.L. currently serves as a scientific advisor to biomodal.

## Acknowledgements

R.Z. was supported by the Vagelos Molecular Life Sciences program at the University of Pennsylvania and C.M.H. by the National Institutes of Health (NIH) T32-GM133398. This work was funded by NIH R01-HG010646 to R.M.K. and F31-HG012892 to C.E.L. We gratefully acknowledge Hao Wu, Julia Leu, Ying Li, and Aleksia Barka for guidance on enzymatic footprinting, Noa Erlitzki on computational analysis, Michael Cory for purfied LexA, and all Kohli lab members for helpful discussions. We thank Eleanor Daniels and Jacob Seiple for technical assistance.

