## Supporting Information for "Mapping Protein Occupancy on DNA with an Unnatural Cytosine Modification in Bio-orthogonal Contexts"

**MATERIALS AND METHODS** ........................................................................ p.S2

**Supplementary References** ........................................................................ p.S7

**SUPPLEMENTARY TABLES AND FIGURES**

**Table S1.** Sequences of oligonucleotides ........................................................p.S8

**Figure S1.** Structural overlay of candidate MTases ........................................p.S9

**Figure S2.** SDS-PAGE of purified M.CviPI variants........................................p.S9

**Figure S3.** Base resolution analysis of λ phage genomic DNA modification.................p.S10

**Figure S4.** GpC-specific CxMTase sequence preference characterization .....................p.S11

**Figure S5.** LexA binding assay ........................................................................p.S12

**Figure S6.** MTase footprinting validation for LexA binding ..........................................p.S13

### MATERIALS AND METHODS

#### Cloning and Design of WT and mutant DNA MTases

For the CpG-specific M.MpeI WT and N374K, the plasmid construct pMG81-M.MpeI-His was used as described.[1] For the GpC-specific and CpC-specific MTases, the pBAD-His-MTase-ZZ-strep (MTase = M.CviPI or M.CviQIX(NdeI15)) constructs were obtained from Addgene (Plasmids #198355 and #198357).[2] Given limited initial evidence of *in vivo* activity with screening of these two plasmids, the expression plasmids were revised to encode pBAD-His-MBP-M.CviPI and pBAD-His-MBP-M.CviQIX(NdeI15), constructed using Gibson assembly. Of note, the 15 amino acid deletion in M.CviQIX(NdeI15) found to be important for solubility was maintained in the M.CviQIX construct.[2]

For mutant construction, the residues homologous to N374 in M.MpeI were identified by aligning the M.MpeI crystal structure (PDB:4DKJ)[3] with AlphaFold3-predicted structures of the tagged constructs bound to appropriate DNA sequences for each: CCACATG5mCGCTGAA for His-MBP-M.CviPI or CCACATG5mCCCTGAA for His-MBP-M.CviQIX(NdeI15) (DNA target sequence contexts underlined, 5mC = 5-methylcytosine).[4] Using the predicted and aligned structures, the amino acid residue with shortest distance relative to Cα of N374 in M.MpeI were identified as Y205 in M.CviPI and N218 in M.CviQIX. For each MTase, K and R mutants of these residues were obtained by performing site-directed mutagenesis via a one-piece Gibson assembly with the variant plasmid confirmed by whole plasmid nanopore sequencing (Plasmidsaurus).

#### *In vivo* activity assessment on *E. coli* gDNA

Activity assessments were performed in *E. coli* using plasmids encoding the WT CpG-specific MTase M.MpeI, variants of GpC-specific MTase M.CviPI, and variants of CpC-specific MTase M.CviQIX(NdeI15). Each construct was transformed into T7 Express (NEB), a $McrA^-$, $McrBC^-$, $EcoBr^-m^-$, $Mrr^-$ cell line that does not restrict methylated DNA. Isolated single colonies were inoculated to 5 mL LB with 100 μg/mL carbenicillin and grown at 37 °C overnight. The saturated overnight culture was diluted ~1:50 into 5 mL of LB with 100 μg/mL carbenicillin. The cultures were grown at 37 °C for 3 hours, moved to 16 °C for 1 hour, and then induced using 5 μL of 30% (w/v) L-arabinose, prior to overnight incubation at 16 °C. 1.5 mL of the overnight cultures was harvested and gDNA was extracted (DNeasy Blood & Tissue Kit, Qiagen) into 50 μL of nuclease-free water and quantified using a nanodrop spectrophotometer. To assess the extent of modification, 200 ng of gDNA was digested using the methylation-sensitive restriction enzyme HaeIII (NEB) at 37 °C for 30 minutes followed by heat inactivation at 80 °C for 20 minutes. The digested DNA were separated via gel electrophoresis (0.8% agarose-TAE) and visualized with SYBR Safe DNA Gel Stain (ThermoFisher).

#### Protein Expression and Purification

Expression of WT and N374K M.MpeI were performed as previously described.[1] Expression of M.CviPI variants (WT, Y205K, Y205R) were performed in T7 Express cells. After transformation, single colonies were inoculated into 100 mL LB with 100 μg/mL carbenicillin and grown overnight at 37 °C. 25 mL of the saturated overnight culture were used to inoculate 1.5 L of LB with 100 µg/mL carbenicillin in baffled flasks. The cultures were grown at 37 °C for 3 hours, moved to 16 °C for 1 hour, and then induced with 3 g/L of L-arabinose, prior to overnight incubation with agitation at 16 °C. Cells were harvested via centrifugation at 7,000 xg for 30 minutes at 4 °C, and pellets were stored in -80 °C. For purification, the pellets were thawed and resuspended in 300 mL Buffer A (20 mM sodium phosphate, pH 7.5, 800 mM NaCl, 10 mM imidazole, and 10% glycerol (v/v)) with 6 cOmplete proteinase inhibitor cocktail tablets (Roche), lysed using a microfluidizer, and centrifuged at 30,000 xg for 30 minutes at 4 °C. The supernatant was passed through a 0.22 µm filter before loading on a gravity column with 3 mL HisPur Cobalt Resin (Thermo Scientific), pre-equilibrated in Buffer A. After loading, the column was washed with at least 20 column volumes (CV) of Buffer A. Elution was performed in three steps, each with 1 CV of Buffer B (20 mM sodium phosphate, pH 7.5, 800 mM NaCl, 200 mM imidazole, and 10% glycerol (v/v)). The purified fractions were pooled, dialyzed (10K MWCO, ThermoFisher) into Storage Buffer (20 mM sodium phosphate, pH 7.5 at 25 °C, 200 mM NaCl, and 10% glycerol (v/v)), and stored at -80°C prior to use. All fractions were were visualized via SDS-PAGE with Coomassie stain and quantified by Qubit Protein Assay (ThermoFisher).

***In vitro* activity assessment on plasmid DNA**

Carboxy-*S*-adenosyl-L-methionine (CxSAM) was synthesized as previously described.[5] Activity assays were performed on 200 ng of unmodified pUC19, using M.CviPI variants, starting at a final enzyme concentration of 780 nM and serially diluted 4-fold (to 12.2 nM), incubated with either 300 µM *S*-adenosyl-L-methionine (SAM) or CxSAM. Reactions were carried out at 37°C overnight in MTase Reaction Buffer (10 mM Tris-HCl, pH 7.9 at 25 °C, 50 mM NaCl, 10 mM EDTA, 1 mM DTT, 4% glycerol, fresh addition of 10 mM TCEP), supplemented with 1 µL of 1 mg/mL BSA (NEB) in a total volume of 10 µL. To help release the MTase from the DNA, the reactions were snap frozen at -80°C for at least 10 minutes prior to purification using Oligo Clean & Concentrator Kit (Zymo) per manufacturer instructions. The product DNA was eluted in 10 µL of nuclease-free water. Purified plasmids were digested for 1 hour at 37 ºC with ApoI (NEB) to linearize all plasmids and with the methylation-sensitive restriction enzyme HaeIII (NEB) to detect modifications, followed by heat inactivation at 80 °C for 20 minutes. The digested DNA products were separated via gel electrophoresis (1% agarose-TAE) and visualized with SYBR Safe DNA Gel Stain (Thermo Fisher).

***In vitro* activity assessment on oligonucleotide duplex**

Activity assays on oligonucleotide substrates were performed with FAM-S27 annealed with S27-Comp-5mC (see **Table S1** for sequences). 100 nM annealed duplex in MTase Reaction Buffer was incubated for >1 hour at 37 ºC with 780 nM M.CviPI WT or mutants and 300 µM SAM or CxSAM, prior to heat inactivation at 95°C for 5 minutes. The reacted samples were then treated with Proteinase K (0.8 Units) or 15 minutes at 37 ºC and heat inactivated at 95 °C for 10 minutes. After snap freezing and storage at -80°C for at least 10 minutes, the oligonucleotides were purified using Oligo Clean & Concentrator Kit (Zymo) and eluted into 6 µL of nuclease-free water. One part of the sample was analyzed by intact oligonucleotide LC/MS (Novatia). For the remaining sample, the complementary strand was exchanged by adding excess S27-Comp-C containing an unmodified C, heating to 95 °C, and slow cooling. The resulting samples were then subjected to restriction digestion by HaeIII at 37 °C for 1 hour followed by heat inactivation at 80 °C for 20 minutes. The reaction products with mixed with loading dye, separated on a 20% acrylamide denaturing gel containing 7 M urea, and imaged using the FAM signal on an Amersham Typhoon.

**Activity assessment on complex genomic DNA substrate**

100 ng of λ phage gDNA (dam$^-$/dcm$^-$, Thermo Fisher) in 20 µL total volume was incubated with 780 nM M.CviPI variants or M.MpeI N374K and 300 µM SAM or CxSAM in MTase reaction buffer. After reaction at 37 ºC overnight, samples were treated with Proteinase K (0.8 Units) at 37 °C for 15 minutes and heat inactivated at 95 °C for 10 minutes. After snap freezing and storage at -80°C for at least 10 minutes, the gDNA samples were then purified using DNeasy Blood & Tissue Kit (Qiagen), eluted in 26 µL of nuclease-free water, and quantified using Qubit dsDNA High Sensitivity (HS) Assay (ThermoFisher). For experiments examining CxMTase activity at higher CxSAM concentration, M.CviPI Y205K was incubated with 800 µM CxSAM and taken through the same workflow.

For analysis of modification levels, the resulting product DNA was used as input into either BS-Seq or ACE-Seq pipelines, with 4 ng or 1 ng of input gDNA, respectively. For both pipelines, the gDNA samples were first sheared to 350 bp using a Covaris sonicator. The sheared DNA was end-repaired with the NEBNext Ultra II End Prep Kit (NEB) and ligated with either IDT xGen Y-shaped adaptors containing either all 5mC for BS-Seq or all 5pyC (custom synthesis from IDT) for ACE-Seq, as previously described.[6] The adaptor-ligated DNA was purified by 1.2X SPRIselect beads (Beckman Coulter) and then quantified by Qubit dsDNA HS Assay (ThermoFisher). BS-Seq was performed with a Premium Bisulfite kit (Diagenode) as per manufacturer instructions. ACE-Seq was performed per the established protocol using reagents from the Enzymatic Methyl-seq Conversion Module (NEB).[7] For library generation, samples were amplified with the Kapa HiFi HotStart Uracil+ReadyMix Kit for 12 cycles. The resulting barcoded libraries were characterized by TapeStation, pooled, and 0.8X left-sided SPRI purified before diluting to 2 nM final library. Sequencing was performed using an Illumina MiSeq with 150 bp paired end reads, with analytical pipeline described below.

**LexA binding and footprinting assays**

To establish if the GpC-specific MTase or CxMTase can be employed to decipher protein-bound and protein-unbound DNA, duplex oligonucleotides were used that contain an embedded LexA binding site, along with a HhaI restriction site. The SOS-box and scrambled DNA sequences (see **Table S1**) were generated by amplification of template DNA using Q5 HotStart High-Fidelity MasterMix (NEB). Amplifications were performed with a reverse primer and either AlexF488 forward binding primer or unlabeled forward binding primer to yield a fluorescent labeled (AlexFluor488) or unlabeled version of each duplex. Products were separated on 2% TAE agarose gel and visualized with SYBR Safe DNA Gel Stain (Thermo Fisher), purified with Oligo Clean & Concentrator Kit (Zymo), and quantified by Nanodrop.

For confirmation of LexA binding, 30 nM of the fluorescent labeled duplex amplicon of interest was incubated with LexA S119A, starting at 780 nM and serially diluted 2-fold (to 94 nM), at room temperature (25 °C) for 20 minutes in LexA binding buffer (10 mM Tris-HCl, pH 7.6 at 25°C, 50 mM NaCl,10 mM EDTA, 10 mM TCEP, 0.04 mg/mL BSA, 5% glycerol) in a 10 µL total reaction volume. The resulting DNA and protein-DNA complex in the product were separated in the gel-shift assay. The total 10 µL resulting product was mixed with loading dye, separated on an 8% native PAGE TBE gel, and visualized for the AlexFluor488-labeled DNA on an Amersham Typhoon imager.

For digestion-based validation of footprinting, we employed conditions identical to those above scaled to 20 µL total reaction volume to evaluate binding, with the addition of 900 nM M.CviPI (WT or Y205K) and either SAM or CxSAM, respectively, at 160 µM. The MTase or CxMTase reactions were performed at 37 °C for 15 minutes and heat inactivated at 95 °C for 5 minutes. The samples were then treated with Proteinase K (0.8 Units) at 37 °C for 15 minutes and heat inactivated at 95 °C for 10 minutes. After snap freezing and storage at -80°C for at least 10 minutes, the samples were purified using an Oligo Clean & Concentrator Kit (Zymo) and eluted in 11 µL of nuclease-free water. 5 µL of the purified product was digested overnight at 37 ºC with the methylation-sensitive restriction enzyme HhaI (NEB) to detect modifications, followed by heat inactivation at 65 °C for 20 minutes. The digested samples were separated on a 20% acrylamide denaturing gel containing 7 M urea and imaged for the FAM signal using an Amersham Typhoon.

For sequencing-based validation of footprinting, we employed binding, footprinting, and clean-up reaction conditions identical to those above with 20 µL total reaction volume. The products were purified using an Oligo Clean & Concentrator Kit (Zymo), eluted in 16 µL, and quantified by Qubit dsDNA HS Assay (ThermoFisher). The resulting DNA samples were then processed using the BS-Seq pipeline described for the λ phage gDNA, with the exception that no shearing was required.

For the experiments with the *lexA* operator, the relevant sequences LexA-Dcm-unmodified or LexA-Dcm-modified duplex, which are with or without 5mC in the Dcm site, respectively, were synthesized as Ultramer duplexes (IDT). The binding, footprinting, and clean-up reaction conditions were identical to those above with 20 µL total reaction volume, except using a final concentration of 300 µM CxSAM. Purification of products and BS-Seq was performed as in sequencing-based validation of footprinting.

**Data analysis**

Sequencing data and code used for processing and analysis used in this study are available at https://github.com/JojoZhu1030/GpC_CxMTase. In general, initial processing (trimming, alignment, deduplication) of raw sequencing reads were done using the protocol described previously.[6] Bismark methylation extractor was run with option CX to obtain information of Cs in all contexts, and the output Bismark coverage files were used for further processing.

For λ phage gDNA samples, the global modification percentage is measured by % reads as C (number of reads as C / total reads at C sites in the reference genome). For % reads as C in GpC context, the Bismark coverage files were filtered in R to isolate sites in GpC contexts, from which the number of reads as C and total covered C were extracted. For analyzing off-target activity in WpCpG contexts, a standard metric for NOMe-Seq data processing given the GpC and CpC activity of M.CviPI WT with SAM, the coverage2cytosine function was run to obtain a WCG report file. The resulting file was processed as for GpC sites to calculate the % reads as C. For each condition, mean and standard deviation were computed across three independent biological replicates (n = 3) using base R functions. One-way ANOVA and Tukey's HSD post-hoc test was done for significance testing. For modification logo, the Bismark coverage files containing Cs in all contexts were used as input. Cytosine sites with coverage <3 times were excluded, and a 4-bp sequence window around the target cytosine were extracted from the reference genome based on the recorded coordinates. Average modification values were calculated (total methylated reads / total reads) and normalized to probability of modification. The sequence logo was made using the ggseqlogo R package.

In read-level analysis, Pearson correlation was performed between % reads as C in GpC sites assessed by BS-Seq and those from ACE-Seq using the cor function in base R. For asymmetry analysis, if the GpC site on either the Watson or Crick strand was covered <15 times the site was excluded from analysis. To avoid overplotting, the data was randomly downsampled to 1000 GpC dyads. Asymmetry was calculated as the absolute value of the difference between the modification percentage on each strand. For sequence context preference analysis around the target site, the data was similarly downsampled to 2000 GpC sites. The sequence context was extracted from the trinucleotide context column in the filtered Bismark coverage file containing only sites in GpC contexts. The points are gittered for better visualization.

For LexA footprinting assays, the reference sequences used for alignmnent were padded with 5 bases of N at both ends to prevent alignment failure. The position reported appropriately adjusted to get the actual position in the sequence. DNA near the ends of linear strands (5 bp) excluded from the GpC modification to avoid confounding from less reliable reads at the end of molecules.

### SUPPLEMENTARY TABLES AND FIGURES

### Table S1. Sequences of oligonucleotides.

| **Name** | **Sequence (5´→3´)** | **Purpose** |
|---|---|---|
| FAM-S27 | FAM-CTATCTAGTTCCATGGCCTCTATAAGA | *In vitro* activity assessment on oligonucleotide duplex |
| S27-Comp-5mC | TCTTATAGAGG5mCCATGGAACTAGATAG | *In vitro* activity assessment on oligonucleotide duplex |
| S27-Comp-C | TCTTATAGAGGCCATGGAACTAGATAG | *In vitro* activity assessment on oligonucleotide duplex |
| SOS-box DNA | ATTGCAGACCTTGTGGCAACAATTTCTAGATGCCTGCG GATGCTGTATGCGCATACAGCATCAATTCTGGCTCAGA GCATATTGACTATCCGGTATTACC | LexA S119A binding assay |
| Scrambled DNA | ATTGCAGACCTTGTGGCAACAATTTCTAGATGCCTGCG GATGCAGTGTGCGCAACATGCATCAATTCTGGCTCAGA GCATATTGACTATCCGGTATTACC | LexA S119A binding assay |
| AlexF488 forward primer | AlexF488-ATTGCAGACCTTGTGGCAACAA | LexA S119A binding assay |
| Forward primer | ATTGCAGACCTTGTGGCAACAA | LexA S119A binding assay |
| Reverse primer | GGTAATACCGGATAGTCAATATGCT | LexA S119A binding assay |
| LexA-Dcm-unmodified DNA | GGTTCCAAATCGCCTTTTGCTGTATATACTCACAGCATAA CTGTATATACACCCAGGGGGCGGAATGAAAGCGTTAACG GC | LexA Dcm footprinting assay (with unmethylated complement) |
| LexA-Dcm-modified DNA | GGTTCCAAATCGCCTTTTGCTGTATATACTCACAGCATAA CTGTATATACACC5mCAGGGGGCGGAATGAAAGCGTTAA CGGC | LexA Dcm footprinting assay (with methylated complement) |

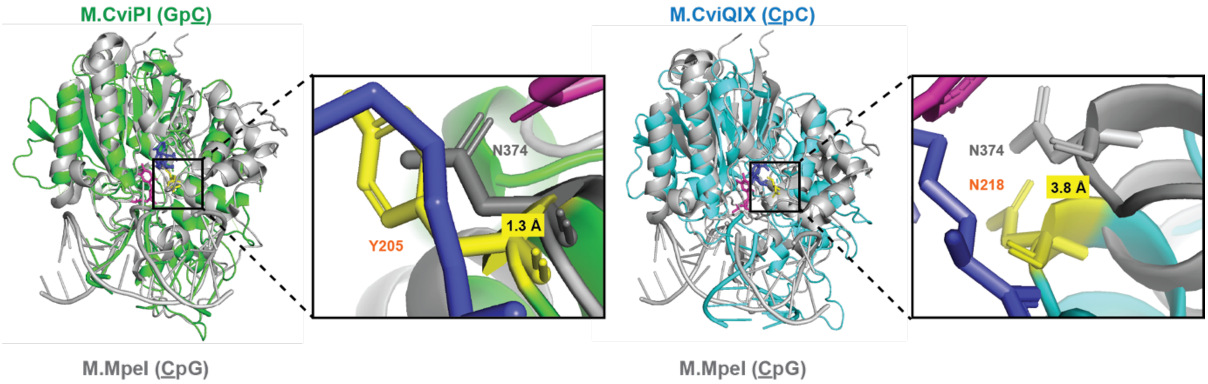


**Figure S1. Structural overlay of candidate MTases.** At left is the structural overlay of an AlphaFold3-predicted structure of M.CviPI (green) bound to Gp5mC-containing DNA with the crystal structure of M.MpeI (grey) bound to 5mCpG-containing DNA and SAH (blue) (PDB 4DKJ). At right is the structural overlay of an AlphaFold3-predicted structure of M.CviQIX(Ndel15) (cyan) bound to 5mCpC-containing DNA with the crystal structure of M.MpeI (grey) bound to 5mCpG-containing DNA and SAH (blue) (PDB 4DKJ). The zoom-in views highlight the region surrounding N374 in M.MpeI with distances from Cα of N374 in M.MpeI to Cα of selected residues (yellow) in the two models labeled (1.3 Å for Y205 of M.CviPI and 3.8 Å for N218 of M.CviQIX(Ndel15)).

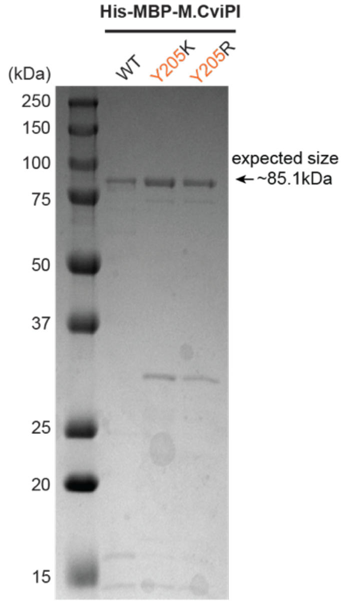


**Figure S2. SDS-PAGE of purified M.CviPI variants.** M.CviPI variants were expressed with an N-terminal His-MBP-tag and purified. The expected size of the full construct is around 85.1 kDa. Purified constructs are shown separated on an SDS PAGE gel.

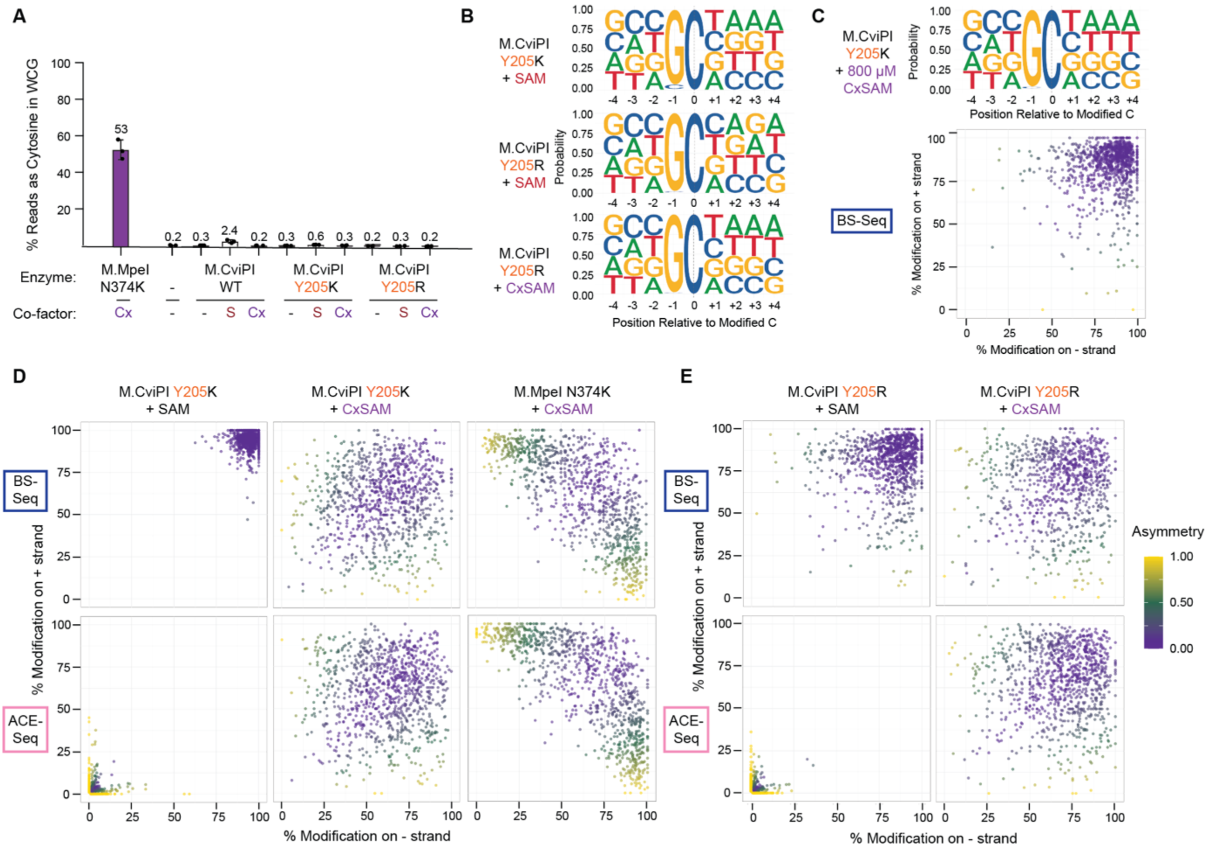


**Figure S3. Base resolution analysis of λ phage genomic DNA modification.** (A) Modification measured by protection of GpC sites from C-to-T conversion (% reads as C) in W<u>C</u>G contexts. G<u>C</u>G and C<u>C</u>G are excluded as in standard NOMe-Seq processing due to overlap with exogenous modification either by on-target or off-target effects. (B) Modification logos of different enzyme and co-factor pairs. The numbering is relative to the target cytosine base (0 position). (C) M.CviPI Y205K off-target and asymmetric carboxymethylation assessment with 800 µM CxSAM. Modification logo and a scatter plot showing the relationship between GpC modifications on opposite strands within the same dyad. Data is filtered for GpCs with at least 15 sequencing reads. Asymmetry is calculated as (% modification on + strand subtracted from % modification on – strand) divided by (sum of + and – strand modification). (D) Scatter plots showing asymmetry of M.CviPI Y205K compared to M.MpeI N374K as assessed by BS-Seq and ACE-Seq. (E) Scatter plots showing asymmetry of M.CviPI Y205R as assessed by BS-Seq and ACE-Seq.

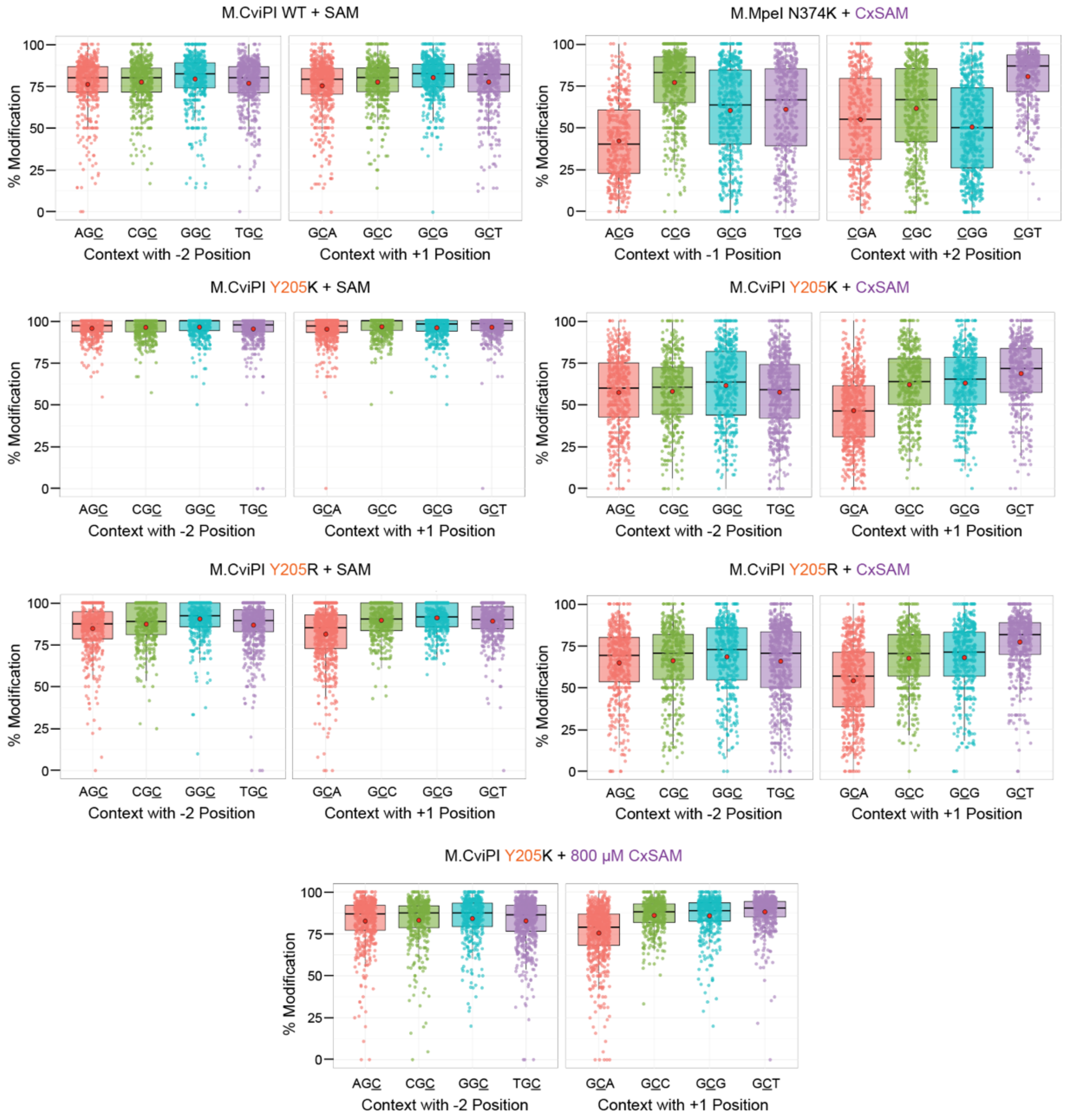


**Figure S4. GpC-specific CxMTase sequence preference characterization.** Modification percentage in target modification context with respect to different bases at immediately adjacent positions. For each enzyme, the numbering is relative to the target cytosine base (0 position). For M.MpeI, analysis includes the -1 position and +2 position identities relative to the CpG target. For M.CviPI, analysis includes the -2 position and the +1 position identities relative to the GpC target.

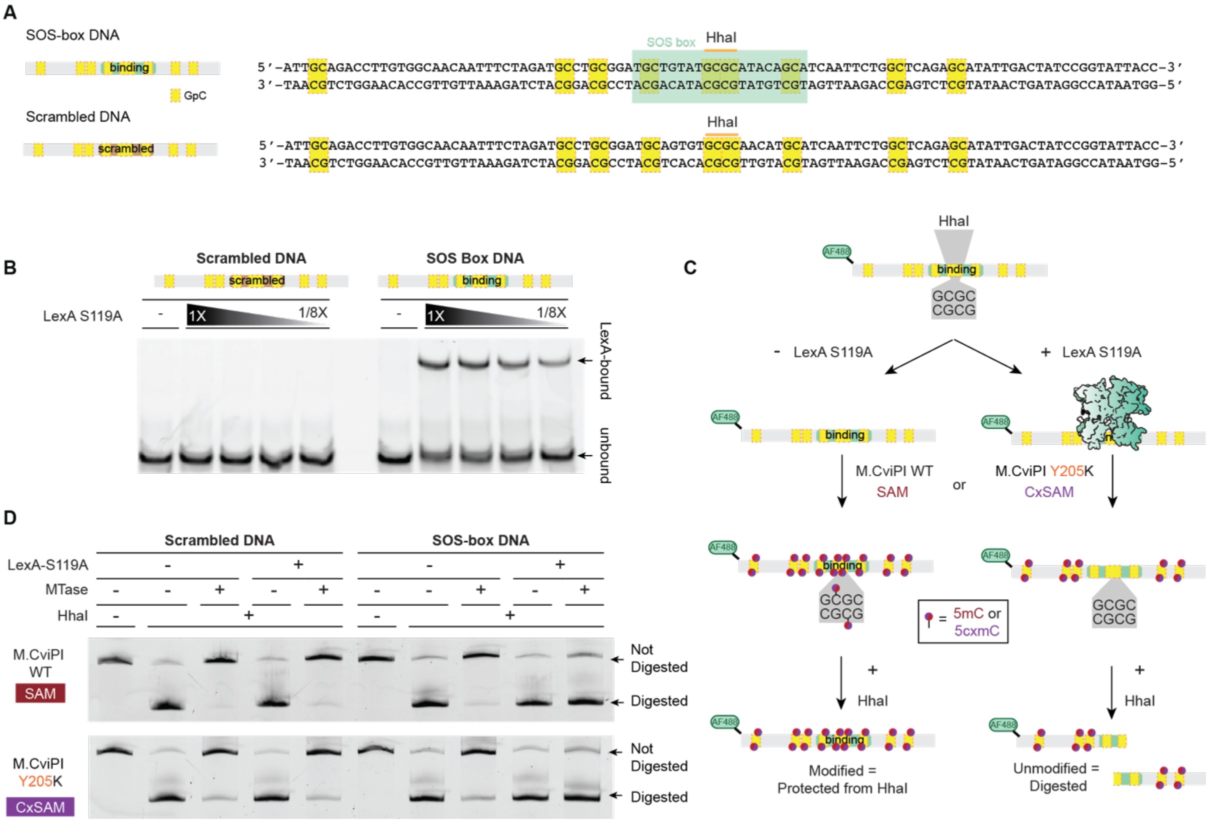


**Figure S5. LexA binding assay.** (A) Scheme of SOS-box and scrambled DNA substrate used. The GpC sites were highlighted in yellow. The SOS box shaded in green with the embedded HhaI site noted (GCGC). The scrambled SOS box is denoted with brown in the schematic at left. (B) Native PAGE of DNA-protein complex after LexA binding. (C) Scheme of LexA binding assay for digestion-based validation of footprinting. SOS-box or scrambled DNA were incubated with catalytically inactive LexA S119A. After the binding reaction, M.CviPI variants and appropriate cofactors (160 µM SAM or CxSAM) were added. The resulting product DNA was evaluated for modification with methylation-sensitive HhaI. (D) Denaturing DNA PAGE of digested samples.

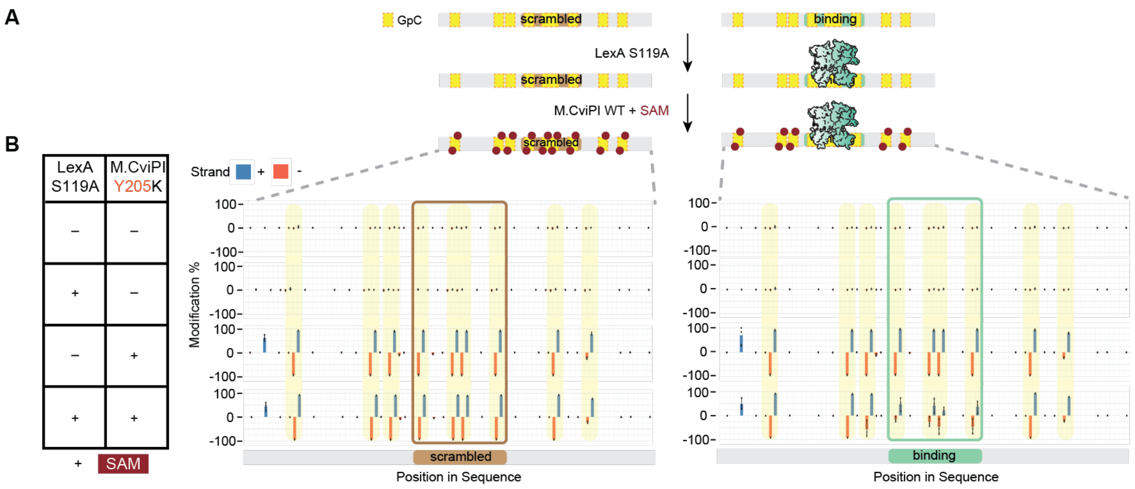


**Figure S6. MTase footprinting validation for LexA binding.** (A) Scheme of LexA footprinting validation assay with WT M.CviPI and SAM. The scrambled 20-bp LexA binding site is indicated by the brown colored box. The 20-bp LexA binding site is indicated by the green colored box. SOS-box or scrambled DNA were incubated with catalytically inactive LexA S119A. After the binding reaction, M.CviPI variants and appropriate cofactors (160 µM SAM or CxSAM) were added. The product DNA was isolated and subjected to BS-Seq to detect modifications at base resolution. (B) Footprinting validation for LexA binding at substrates containing either a scrambled or unscrambled binding site with WT M.CviPI and SAM. Modification level on + strand (blue, upward) and – strand (orange, downward) assessed by BS-Seq. Shown are individual data points from independent replicates (n = 3), with mean values in bars and standard deviation noted.
